# Where have all the spiders gone? Observations of a dramatic population density decline in the once very abundant garden spider, *Araneus diadematus* (Araneae: Araneidae), in the Swiss midland

**DOI:** 10.1101/2020.03.03.975912

**Authors:** Martin Nyffeler, Dries Bonte

## Abstract

Aerial web-spinning spiders (including large orb-weavers) depend, as a group of insectivores, completely on flying insects as a food source. The recent widespread loss of flying insects across large parts of western Europe, both in terms of diversity and biomass, can therefore be anticipated to have a drastic negative impact on survival and abundance of this type of spiders. To test the putative importance of such a to date neglected trophic cascade, a survey of population densities of the European garden spider *Araneus diadematus* – a large orb-weaving spider – was conducted in late summer 2019 on twenty sites of the Swiss midland. The data from this survey were compared with published population densities for this species from the previous century. The study verifies above-mentioned hypothesis that this spider’s present-day overall mean population density has declined alarmly to densities much lower than can be expected from normal population fluctuations (0.7% of the historical densities). Review of other available records suggests this pattern is widespread and not restricted to this region. In conclusion, the here documented abundance decline of this once so abundant spider in the Swiss midland is evidently revealing a bottom-up trophic cascade in response to the widespread loss of flying insect prey in recent decades.

## 1. Introduction

When the Entomological Group Krefeld along with colleagues published their famous long-term study (the “Krefeld study”) in 2017, it became known that the flying insect biomass had declined by approx. 75% over the past three decades in over 60 nature protection areas of Germany [1]. Another long-term study (the “Munich study”) basically confirmed the results of the Krefeld study, providing evidence that strong abundance declines of insect populations had occurred in farmland and forests across vast areas of Germany and Switzerland [2]. Similar abundance declines of insect populations were documented by means of other long-term studies for other European and North American regions [3–9]. Meanwhile, it is generally accepted as scientifically proven that in many parts of the globe insect population densities had drastically decreased in recent decades and that we now live in an era of global insect meltdown [9–13]. This has dramatic ecological implications due to the fact that insects comprise the base of many food chains and food webs [14,15] – in a world without insects countless insectivorous species would ultimately become extinct due to starvation [16].

In this context, aerial web-spinning spiders are a group of uttermost interest. This group of spiders (including the orb-weaving spiders in the families Araneidae) trap flying insects with the aid of aerial webs [17–19]. In the temperate climate zone, aerial web-spinning spiders feed for the most part on small dipterans, aphids, and hymenopterans (Figure 1) which are exactly the type of insects known to have most dramatically decreased in abundance and biomass in recent decades [1]. With other words, because of their reliance upon flying insects as their principal food source, aerial web-spinning spiders are a highly vulnerable predator group in regions characterized by high loss of flying insects such as Germany and Switzerland.

**Figure 1.**
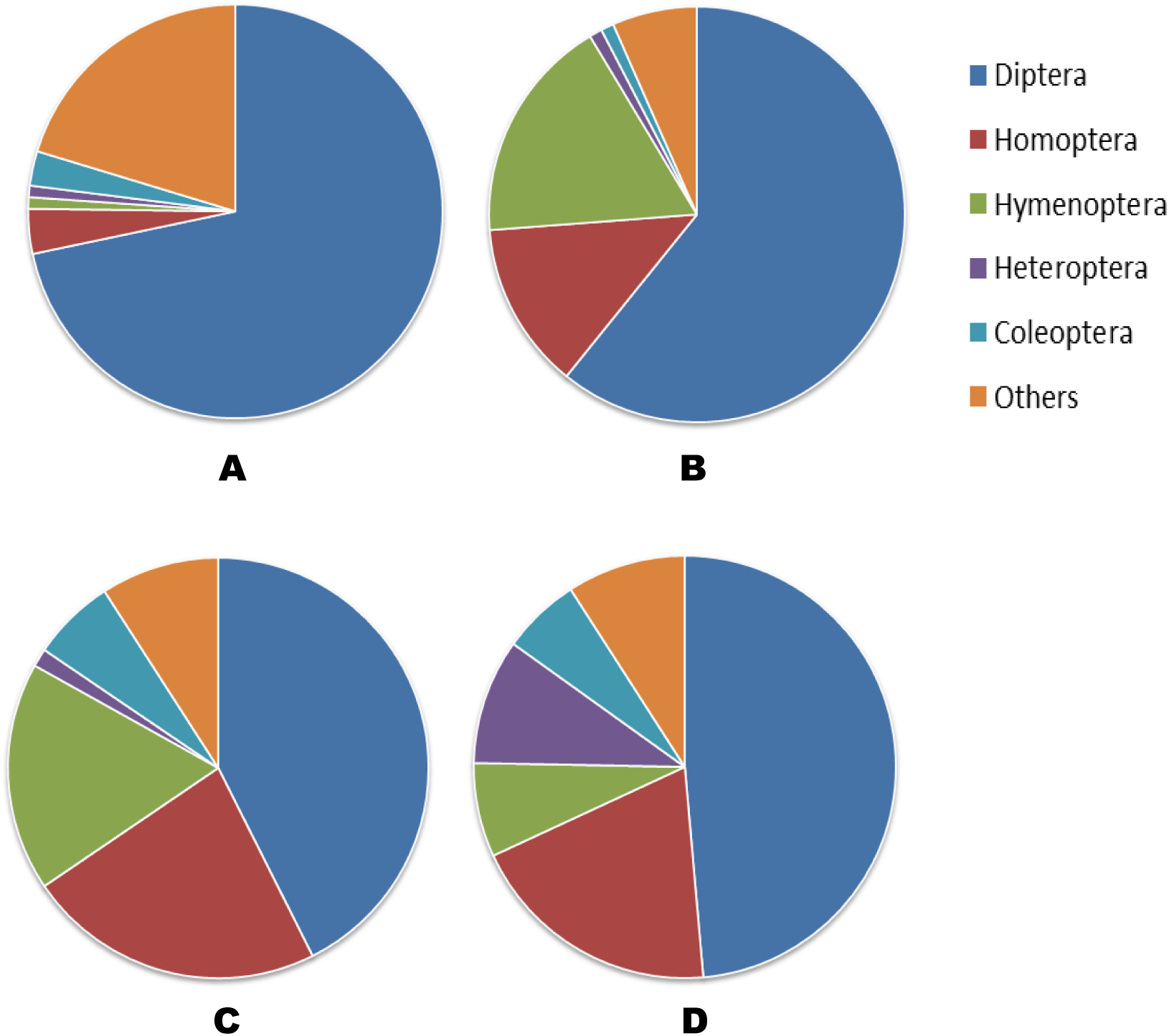
Prey composition (% by number) of the garden orb-weaver *Araneus diadematus* based on four field studies. A) Oat field near Zurich, Switzerland [21]. B) Fallow grassland near Zurich, Switzerland [21]. C) Fallow grassland near Jena, Germany [18]. D) Maize field margin near Munich [19]. Homoptera refers primarily to alate aphids.

While a larger number of studies on population declines of various groups of insects had been conducted in recent years (see references in [12]), potential cascading effects have been documented so far for insectivorous birds [6]. No studies of this type have so far considered potential declines due to trophic cascades among arthropods, like aerial web-spinning spiders. In the following we use the European garden spider *Araneus diadematus* as a model system to address the question whether evidence for population declines in aerial web-spinning spiders in the Swiss midland can be found. The female, 10–18 mm in length, reaches adulthood in late summer, at which time it spins (≈30 cm Ø) orbs [20–22]. The webs are built 0.5–2.5 m off the ground [21]. This animal with its conspicuous white, cross-shaped mark on the upper side of the abdomen is one of the best known spider species in western Europe.

By comparing historical abundance data (20^th^ century) with present-day data (year 2019) it shall be examined whether the abundance of *Araneus diadematus* has changed over the past few decades. This work is based on extensive experience gathered during graduate research in the Swiss midland in the 20^th^ century [20,21] and has since been supplemented by a present-day population density survey in the same geographical area. To make the comparison more robust, the data base was expanded by adding published population density values extracted from the scientific literature.

## 2. Materials and Methods

### 2.1. Assessment of population densities of Araneus diadematus in 2019

Population density assessments were conducted on 20 locations in the Swiss midland by walking transects and counting webs (see Table 1, Table S1 for details). The search focussed on webs of adult females in its typical habitats (gardens, parks, graveyards, hedgerows, forest edges, forest trails, fallow grassland patches). Transects of 200 to 1000 m^2^ ground area were used (area size depending upon the specific situation; see Table S1 for details). All counts were conducted on rain-free afternoons in August and September 2019.

**Table 1.** Population densities of adult female *Araneus diadematus* (18 historical vs. 20 recent values). *Adults/subadults in September

| Habitat type | Geographic region | Period of investigation | Spiders m<sup>-2</sup> | Source |
| --- | --- | --- | --- | --- |
| **<u>Historical data\*</u>** |  |  |  |  |
| Fallow grassland, plot 1 | Switzerland | 1979 | 0.000 | [21] |
| Fallow grassland, plot 2 | Switzerland | 1979 | 0.030 | [21] |
| Fallow grassland, plot 3 | Switzerland | 1979 | 0.070 | [21] |
| Fallow grassland, plot 4 | Switzerland | 1979 | 0.000 | [21] |
| Shrubs | Switzerland | 1979 | 0.930 | [21] |
| Garden | Switzerland | 1974–75 | 0.064 | [20] |
| Glasshouses Botanical Garden Bern | Switzerland | 1932 | 0.001 | [71] |
| Fallow grassland | Germany | 1990 | 0.138 | [18] |
| Fallow grassland | Germany | 1998 | 0.056 | [32] |
| Fallow grassland | France | 1984 | 0.023 | [30] |
| Hedgerow/grassland | France | 1977 | 0.135 | [26] |
| Hedgerow | Italy | 1990–91 | 0.0005 | [31] |
| Pine stand | Poland | 1979 | 0.076 | [28] |
| Clearing-forest ecotone | Poland | 1979 | 0.320 | [28] |
| Scotch pine wood | Netherland | 1946 | 0.130 | [29] |
| Oak wood | Netherland | 1946 | 0.400 | [29] |
| Oak stand | UK | 1954–56 | 0.410 | [72] |
| Heathland | UK | 1968 | 0.024* | [27] |
| **Overall mean ± SE** |  |  | 0.1560 ± 0.0552 |  |
| **<u>Recent data</u>** |  |  |  |  |
| Forest road | Northeastern Switzerl. | 2019 | 0.002 | This paper |
| Forest edge + suburb | Northwestern Switzerl. | 2019 | 0.002 | This paper |
| Forest road | Northwestern Switzerl. | 2019 | 0.000 | This paper |
| Forest edge | Northwestern Switzerl. | 2019 | 0.000 | This paper |
| Forest edge | Northwestern Switzerl. | 2019 | 0.000 | This paper |
| Forest edge | Northwestern Switzerl. | 2019 | 0.000 | This paper |
| Forest road | Northwestern Switzerl. | 2019 | 0.000 | This paper |
| Garden | Northeastern Switzerl. | 2019 | 0.006 | This paper |
| Garden | Northwestern Switzerl. | 2019 | 0.003 | This paper |
| Garden + shrubs | Northwestern Switzerl. | 2019 | 0.000 | This paper |
| Graveyard shrubs | Northwestern Switzerl. | 2019 | 0.005 | This paper |
| Graveyard shrubs | Northwestern Switzerl. | 2019 | 0.000 | This paper |
| Public park | Northwestern Switzerl. | 2019 | 0.000 | This paper |
| Suburb | Zurich region, Switzerl. | 2019 | 0.000 | This paper |
| Suburb | Zurich region, Switzerl. | 2019 | 0.000 | This paper |
| Suburb | Northwestern Switzerl. | 2019 | 0.001 | This paper |
| Hedgerow | Northwestern Switzerl. | 2019 | 0.000 | This paper |
| Hedgerow | Northwestern Switzerl. | 2019 | 0.000 | This paper |
| Hedgerow + reedbelt | Central Switzerl. | 2019 | 0.000 | This paper |
| Hedgerow + shrubs along a river bank | Zurich region, Switzerl. | 2019 | 0.002 | This paper |
| **Overall mean ± SE** |  |  | 0.0011 ± 0.0004 |  |

### 2.1. Literature search for historical data on Araneus diadematus population densities

*Araneus diadematus* population density values assessed in the 1970s have been taken from two graduate research theses which had been conducted in the Zurich area [20,21]. These data were supplemented by data taken from the literature. Overall, a total of 12 reports (including 18 population density values) could be gathered (see Table S1). In these studies, transects ranging from 100 to 5000 m^2^ ground area were used (see Table S1).

### 2.3. Statistical methods

The Mann–Whitney *U* test was applied to examine whether overall mean spider density in 2019 (N = 20) differed statistically significantly from overall mean spider density in the 20^th^ century (N = 18). Mean values are followed by Standard Errors (± SE).

## 3. Results

In Table 1 population densities of *Araneus diadematus* assessed in the Swiss midland in August/September 2019 are compared with 20^th^ century population density data for this species from a variety of European locations. The table reveals that present-day population densities of adult female *Araneus diadematus* in the Swiss midland were generally extremly low (overall mean = 0.0011 ± 0.0004 webs m^-2^ [range = 0.000–0.006 webs m^-2^]; Table 1). In 2/3 of the 20 investigated transects no webs of adult female *Araneus diadematus* could be found.

By contrast, historical population densities of *Araneus diadematus* in its typical habitats had been considerably higher (overall mean = 0.1560 ± 0.0552 webs m^-2^; Table 1. The difference between the 20^th^ century European overall mean and the present-day Swiss overall mean (0.1560 vs. 0.0011 webs m^-2^) is statistically highly significant (Mann–Whitney *U* test, n_1_ = 18, n_2_ = 20, Z = −4.0637, p < 0.001). The present-day Swiss overall mean density of *Araneus diadematus* is roughly 140 times lower than the 20^th^ century European overall mean (Table 1). [If the overall mean of exclusively the seven Swiss 20^th^ century values (0.1564 webs m^-2^, Table 1) is compared with the overall mean of the twenty Swiss 21^th^ century values (0.0011 webs m^-2^, Table 1), the ratio still remains 140 : 1, and the difference between the two overall means is still statistically significant at p < 0.05.]

In 2019, the webs appeared to be rather fragile (Figure 2A–B) compared with the stronger webs from previous decades, as is the case when malnourished spiders make webs with thinner threads (Nyffeler, pers. observations). This is in good agreement with laboratory experiments, in which the amount of produced thread was reduced if *Araneus diadematus* spiders were kept fooddeprived [23]. Furthermore, the webs contained few prey compared to previous studies (Table S2). The overall impression gained during this survey was that there was a paucity of prey throughout the entire study area in recent decades which is confirmed by the observations of other researchers [2,24,25].

**Figure 2A–B.**
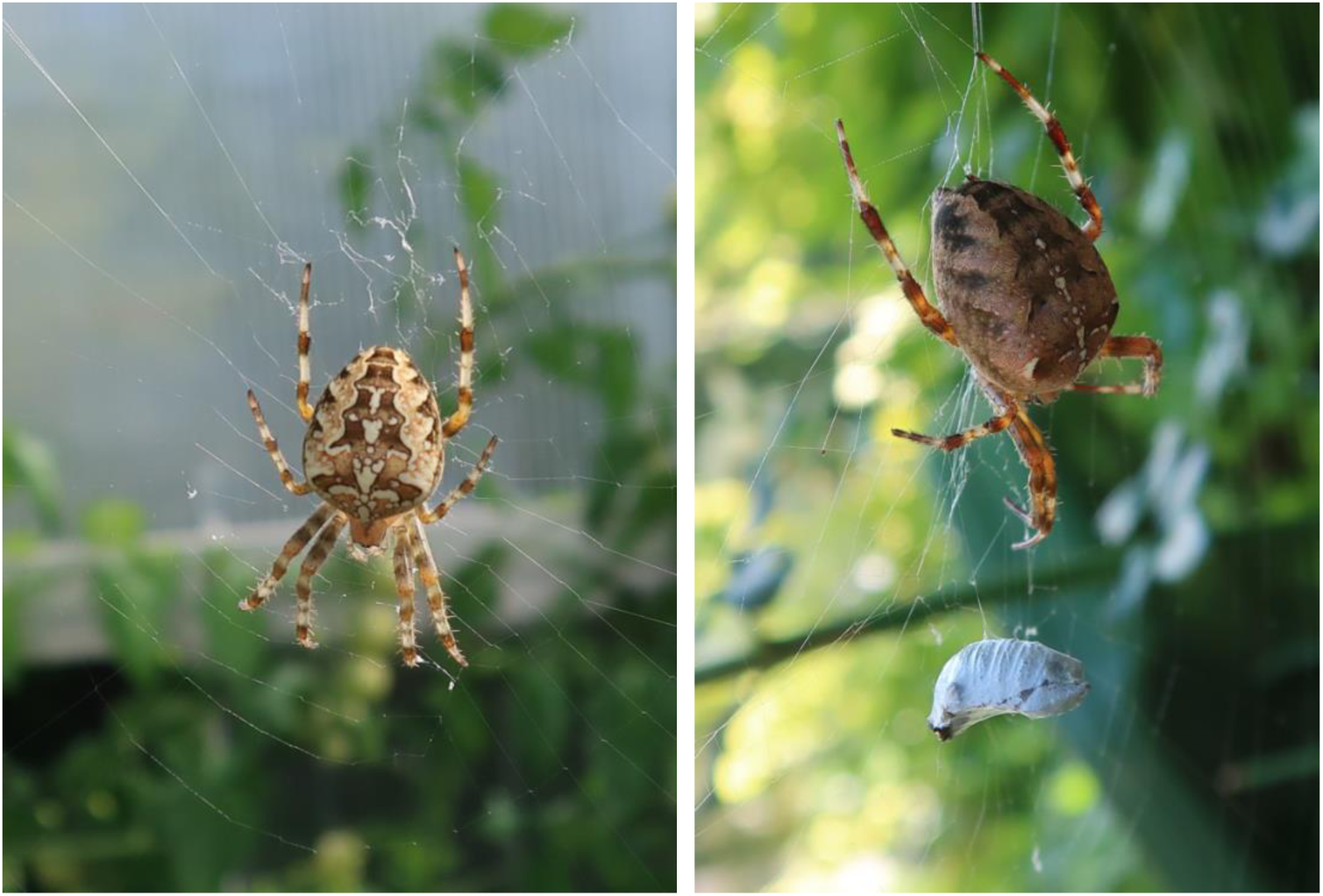
Adult female *Araneus diadematus* in web in a garden in Flawil, northeastern Switzerland (courtesy: Rätus Fischer)

## 4. Discussion

This study revealed that the large orb-weaving spider *Araneus diadematus* occurred in extraordinarily low population densities in the Swiss midland in 2019 (Table 1). *Araneus diadematus* spiders take down their webs during the night and rebuild them early in the morning. However, as laboratory experiments and field observations had revealed, not the entire population is rebuilding its webs the following day [23,26]. Some spiders, which had consumed exceptionally large amounts of food at particularly favorable hunting sites, may cease feeding and rebuilding their webs for one or several days [23]. Thus, the percentage of spiders found with webs within a *A. diadematus* population is in general lower than 100% (usually ranging between about 66–88%) depending on the availability of food at a particular time and location [23,26]. The question arises whether the method of web counts applied in this study may have resulted in underestimating the true population densities by overlooking a certain percentage of satiated spiders which had not rebuilt their webs at the time when the counts were conducted. On this issue, it may be noted that densities assessed in the past century were for the most part also based on web counts [18,20,21,26–32] so that the same methodological bias was involved in both time periods. Furthermore, since insatiated spiders tend to rebuild their webs more frequently than fully satiated spiders [23,33], and since present-day spiders living in the era of insect loss are more likely to be insatiated compared with half a century ago (Table S2), the frequency of web building nowadays can be expected to be rather higher compared with the situation in the last century. Thus, the possibility that the extraordinarily low present-day population densities were underestimations due to this particular type of methodological bias can be ruled out.

The here documented abundance decline of this once so abundant spider in the Swiss midland is evidently revealing a bottom-up trophic cascade in response to prey scarcity recently documented in the Swiss midland and across vast areas of western Europe [1,2,24,25]. The hypothesis that the availability of flying insect prey in the study area had drastically declined over the past decades is confirmed by the “windshield phenomenon” noticed throghout the Swiss midland (i.e., nowadays many times less flying insects are killed on the front windshields of cars compared to previous decades; [9,34]. This is in sharp contrast to the situation a few decades ago, when fairly frequently “wasteful killing” (and coupled with it “partial consumption”) of insects in *Araneus diadematus* webs at favorable web sites could be witnessed (with capture rates of sometimes up to 1000 prey web^-1^ day^-1^; [35]). The reduced food intake of *Araneus diadematus* in recent years (Table S2) is assumed to have negatively impacted the fecundity and survival of this spider [36,37) which in turn led to the abundance decline documented in this study (Table 1). Sublethal effects of chemical pollution may have additionally negatively affected this spider’s survival [38–43].

Aside from food scarcity due to insect population declines and chemical pollution, other negative environmental developments additionally contributed to the decreasing population densities of large orb-weaving spider species in the western European landscape. This includes adverse effects of modern forestry practices (e.g., removal of forest understory and of shrubs at forest margins, large-scale aerial spraying for the purpose of caterpillar or bark beetle control), the transformation of traditional farmland into large-scale depauperate monocultures (accompanied by the deletion of weedy field margins and hedges), the transformation of fallow grassland patches into construction sites, and the removal of shrubs and trees in urban and suburban gardens. That habitat conversion and degradation can have a strong detrimental effect on large orb-weaving spider abundance has impressivly been demonstrated on the example of the fate of a plot of land located near Zurich-Oberenstringen (in between highway A3 and the Limmat riverbank). In the 1970s, tall grassland interspersed with shrubs covered this plot and at that time large orb-weaving spiders occurred there in high numbers (up to 6 m^-2^ in small local patches; [44]). During a visit of this site, 40 years later, it became apparent that the land had been converted to a highway rest area with short-cut lawns and, as a result, the density of large orb-weaving spiders had declined to a very low value of 0.002 m^-2^ (Nyffeler, pers. observations).

We of course realise that the recorded declines were not based on systematic long-term monitoring and that the declines could be attribited to normal population cycling. However, while we are not able to provide a smoking gun herein, several elements support a systematic decline. First of all, the dramatic decline up to <1% of the reference baseline from 40 years ago, has never been observed at shorter temporal time frames, and reach far beyond the typical natural fluctuations in population sizes (e.g., Green 2003), even when they are known to be linked to resource pulses (e.g., Wolf 1996). Second, records from other location show that this decline is occuring at such large spatial scales that can only be explained by large-scale, more global environmental drivers like land-use change, climate change or the global use of pollutants. Indeed, the extraordinarily low population densities in *Araneus diadematus* in the Swiss midland observed in the 2019 survey (Table 1) are validated by statements of biology students of the University of Basel according to which generally very few large orb-weaving spiders had been spotted in recent years during walks through forests and fields in the Basel region. Furthermore, in mid-June 2017, a group of over 40 biology experts conducted an extensive faunistic survey on the grounds of the Merian Gardens, a park-like area covering 180,000 m^2^ located in the outskirts of Basel ([47]). In the course of this survey only three specimens of *A. diadematus* could be found over a time period of 24 hours suggesting that nowadays this once so “abundant garden spider” ([48]) must have become rare in that park-like garden area. The only large orb-weaver species still found in high density during the 2019 survey in the Swiss midland was the bridge spider *Larinioides sclopetarius* (Nyffeler, pers. observations). This nocturnal species, which lives near water and frequently builds its webs on street lights and illuminated bridge railings, gets still sufficient amounts of food in the form of flying adults of semiaquatic insects (chironomids, ephemeropterans, trichopterans, etc.) attracted in large numbers to the artificial light [49–51]. Due to its capability to exploit this type of artificially high prey densities, *L. sclopetarius* is – contrary to other large orb-weavers – even in the 21^th^ century extremely successful in colonizing urban habitats in high density, not only in Switzerland, but also in other parts of western Europe [50].

Similar extraordinarily low population densities of *Araneus diadematus* (0.0004 adult females m^-2^ at a landscape scale) were also recorded in an extensive survey in nine lanscapes of the Ghent region, northern Belgium, in summer of 2014 ([43]). Here as well, densities were one order of magnitude or more lower than those recorded one decade ago (e.g., during a survey in 2004–2006 densities at two locations were 10–20 times higher compared to ten years later; [52]; Bonte, pers. observations). Interestingly, while other orb-weaver species – especially the larger ones – showed dramatic declines in both abundance and species richness along an urbanisation gradient [53,54], *A. diadematus* was found to reach similar low densities across this land-use gradient. Or put conversely, local densities at spatial scales of approximately 100 km were alarmingly low in both the more natural and human impacted regions, demonstrating regional declines beyond the scale of local land-use. Thus, extraordinarily low present-day population densities of *Araneus diadematus* have been recorded in two different regions of western Europe, ≈500 km apart. These declines extend local environmental changes at small scales and suggest common negative impacts of intensive urbanization, climate change or any other large-scale stressor across the entire landscape over the past half-century [55,56].

The present study and that from northern Belgium suggest that the notion that *Araneus diadematus* is an abundant spider, as pointed out in much of the literature [57,58], has these days turned out to be a myth – at least in highly urbanized western European landscapes such as the Swiss midland and northern Belgium. But so far the apparent abundance decline of this species in western Europe had been ignored by faunists in charge of compiling local Red Lists for spiders (e.g., [57,58]). Red Lists are based on the number of different locations within a landscape from which a species has been recorded – and not on absolute population densities, and the fact that *Araneus diadematus* is still found in many places (although in strongly reduced densities) may explain why it is still labeled as an “abundant species” (e.g., [57,58]).

In contrast, present-day population densities of *Araneus diadematus* in some areas outside of continental western Europe appear to be still rather high. So for instance, exterminators of the pest control firm Senske Services removed more than 75 *Araneus diadematus* spiders from the exterior of a “2,500–square-foot home” in Seattle (USA) in August 2016, which translates to a population density of 0.32 spiders m^-2^ [59]. Evidently the spiders at this particular location were still capturing a sufficient amount of food in the form of flying insects enabling them to build up a population density roughly 290 times that of the present-day Swiss overall mean density. Future detailed assessments of the population densities of this spider species in the different parts of its geographic range are urgently needed and may provide important clues on the extent of the loss of flying insects in various regions of western Europe and of the globe. We would in this respect like to promote the use of new citizin-science tools such as *SpiderSpotter* [60] to achieve these highly needed insights, and especially to guide short-term action to mitigate these alarming declines in these sentinel arthropod top-predators.

## 5. Concluding Remarks

The drastic decline in the abundance of the orb-weaving spider *Araneus diadematus* over the past half-century documented in this study (Table 1) is apparently revealing a bottom-up trophic cascade in response to widespread insect losses that had occurred across large parts of Europe in recent decades [1,2,4,6,9,25,61,62]. There is evidence that other groups of aerial web-spinning spiders, which likewise depend on flying insects as food [17–19,21,63–65], have also become much rarer over the past decades (Nyffeler, pers. observations). So for instance, the mesh-web weaver *Dictyna uncinata* (along with other dictynid species) whose small, tangled webs were found in large numbers on the leaves of garden plants a few decades ago [65], has apparently become very rare these days (Nyffeler, pers. observations). The ongoing abundance decline of the spiders parallels the dramatic abundance declines in other insectivorous animals such as insectivorous birds, bats, frogs, and lizards documented in recent decades [66–68].

To sum this up, the findings of this study support the notion by other researchers that we now live in the midst of an ecological crisis in which trophic webs are being eroded and degraded as a result of man-made adverse environmental impacts [13,61,67,69,70]. If this disastrous trend cannot be halted or even reversed in the very near future, “entire ecosystems will collapse due to starvation” [16].

## Supplementary Materials

Table S1 and Table S2 are available online at ………………..

## Author Contributions

Conceptualization, M.N and D.B.; Methodology, M.N.; Validation: M.N. and D.B.; Formal Analysis: M.N. and D.B.; Investigation, M.N. (in Switzerland) and D.B. (in Belgium); Data Curation, M.N.; Writing – Original Draft Preparation, M.N.; Writing – Review & Editing, D.B.

## Acknowledgements

The authors wish to extend their gratitude to Rätus Fischer and Christlinde Ruchti for assistance in the field.

