## Supplementary material for "Where have all the spiders gone? Observations of a dramatic population density decline in the once very abundant garden spider, *Araneus diadematus* (Araneae: Araneidae), in the Swiss midland"

### Table S1. Area size ($m^2$) of the study areas

| Habitat type | Geographic region | Study area ($m^2$) |
|---|---|---|
| **Historical data** | | |
| Fallow grassland, plot 1 | Canton Zurich, Switzerl. | ~1000 |
| Fallow grassland, plot 2 | Canton Zurich, Switzerl. | ~1000 |
| Fallow grassland, plot 3 | Canton Zurich, Switzerl. | ~1000 |
| Fallow grassland, plot 4 | Canton Zurich, Switzerl. | ~1000 |
| Shrubs | Canton Zurich, Switzerl. | ~1000 |
| Organic garden | Canton Zurich, Switzerl. | 450 |
| Glasshouses Botanical Garden Bern | Canton Bern, Switzerl. | 965 |
| Fallow grassland | Germany | 4 x 625 |
| Fallow grassland | Germany | 90 |
| Fallow grassland | France | 3000 |
| Hedgerow/grassland | France | ~400* |
| Hedgerow | Italy | 2000 |
| Pine stand | Poland | 100 |
| Clearing-forest ecotone | Poland | 100 |
| Scotch pine wood | Netherland | 157.5** |
| Oak wood | Netherland | 17.5 |
| Oak stand | UK | 2500 |
| Heathland | UK | 1450 |
| **Present-day data (2019)** | | |
| Forest road, Degersheim | Canton St. Gallen. | 1000 |
| Forest edge + suburb, Arlesheim | Canton Baselland | 500 |
| Forest road, Reinacherheide | Canton Baselland | 200 |
| Forest edge, Münchenstein | Canton Baselland | 500 |
| Forest edge, Bottmingen | Canton Baselland | 200 |
| Forest edge, Rheinach | Canton Baselland | 200 |
| Forest road, Leymen | Alsace, France*** | 500 |
| Organic garden, Flawil | Canton St. Gallen | 800 |
| Organic garden, Himmelried | Canton Solothurn | 330 |
| Organic garden + shrubs, Dornach | Canton Solothurn | 500 |
| Graveyard shrubs, Binningen | Canton Baselland | 200 |
| Graveyard shrubs, Oberwil | Canton Baselland | 500 |
| Public park, Riehen | Canton Basel | 200 |
| Suburb, Hönggerberg | Canton Zurich | 1000 |

| | | |
|---|---|---|
| Suburb, Zürichberg | Canton Zurich | 1000 |
| Suburb, Rheinfelden | Canton Aargau | 1000 |
| Hedgerow, Reinach | Canton Baselland | 200 |
| Hedgerow, Schönenbuch | Canton Baselland | 200 |
| Hedgerow + reedbelt, Sempach | Canton Luzern | 200 |
| Hedgerow + shrubs along a river bank, Oberengstringen | Canton Zurich | 1000 |

Additional information to Table S1

* Roughly estimated based on information (i.e., map of web distribution) found in Le Berre, M.; Ramousse, R.; Le Guelte, L. Eco-éthologie des Argiopidae: 1. Evolution temporelle d'une population d'*Araneus diadematus* Clerck dans son milieu naturel. Atti. Soc. Toscana Sci. Nat. P. V. Mem. Ser. B **1981**, *88*, 72-83.

** 20.5 ind/(9 x 17.5 $m^2$) = 20.5 ind/157.5 $m^2$ = 0.130 ind/$m^2$

*** Location in France, only 1.5 km from the Swiss border (Basel region)

**Table S2. Mean number of prey per web counted in mid-afternoon as a proxy for the daily prey capture rate of large orb-weaving spiders in western European habitats: Historical values vs. present day value. References found in the paper.**

| Spider species | Habitat type | Year of investigation | $\bar{x}$ prey web$^{-1}$ | Reference |
|---|---|---|---|---|
| **Historical data:** | | | | |
| *Araneus diadematus* | Grassland, September | 1979 | 13.5 | Nyffeler 1982 |
| *Araneus diadematus* | Grassland, August | 1990 | 4.3 | Malt 1996 |
| *Araneus diadematus* | Grassland, September | 1990 | 9.2 | Malt 1996 |
| *Araneus diadematus* | Maize field margins, July/August | 2003 | 12.5 | Ludy 2007 |
| *Araneus quadratus* | Grassland, July | 1979 | 21 | Nyffeler 1982 |
| *Araneus quadratus* | Grassland, August | 1979 | 14 | Nyffeler 1982 |
| *Araneus quadratus* | Grassland, September | 1979 | 20 | Nyffeler 1982 |
| *Araneus quadratus* | Grassland, August | 1990 | 7.3 | Malt 1996 |
| *Araneus quadratus* | Grassland, September | 1990 | 36.4 | Malt 1996 |
| *Araneus marmoreus* | Grassland | <1984 | 14.1 | Pasquet 1984 |
| *Larinioides cornutus* | Grassland, June/July | 1979 | 18 | Nyffeler 1982 |
| *Larinioides cornutus* | Rye field | 1976 | 15 | Nyffeler 1982 |
| **Overall mean ± SE** | | | **15.44 ± 2.37** | |
| **Present-day data:** | | | | |

| | | | |
|---|---|---|---|
| *Araneus diadematus* | Diverse habitats<br>August/September | 2.74 | This paper |
| **Overall mean ± SE** | | **2.74 ± 0.98** | |
